# Quantifying predator dependence in the functional response of generalist predators

**DOI:** 10.1101/082115

**Authors:** Mark Novak, Christopher Wolf, Kyle Coblentz, Isaac Shepard

## Abstract

A longstanding debate concerns whether functional responses are best described by prey-dependent versus ratio-dependent models. Theory suggests that ratio dependence can explain many food web patterns left unexplained by simple prey-dependent models. However, for logistical reasons, ratio dependence and predator dependence more generally have seen infrequent empirical evaluation and then only so in specialist predators, which are rare in nature. Here we develop an approach to simultaneously estimate the prey-specific attack rates and predator-specific interference rates of predators interacting with arbitrary numbers of prey and predator species. We apply the approach to field surveys and two field experiments involving two intertidal whelks and their full suite of potential prey. Our study provides strong evidence for the presence of weak predator dependence that is closer to being prey dependent than ratio dependent over manipulated and natural ranges of species abundances. It also indicates how, for generalist predators, even the qualitative nature of predator dependence can be prey-specific.

**Author contributions:** CW contributed to method development, KC and IS performed the caging experiment, and MN conceived of the study, carried out the fieldwork and analyses, and wrote the manuscript.

## Introduction

How predator feeding rates respond to changes in prey abundance underlies the dynamics of all predator-prey interactions (Murdoch & Oaten, 1975). A longstanding and still vigorous debate in the predator-prey literature concerns whether these functional responses are best described by prey-dependent models, such as the classical Holling type forms, or by ratio-dependent models (Abrams, 2015; Abrams & Ginzburg, 2000; Arditi & Ginzburg, 2012; Barraquand, 2014). In the former, feeding rates respond only to changes in prey abundance. In the latter, feeding rates respond to the prey available per predator. Predator-dependent functional responses more generally encapsulate the hypothesis that predator individuals alter each others feeding rate and include ratio dependence as a special case. Models including predator dependence indicate that the effective refuge that prey experience at high predator-to-prey ratios can explain many of nature’s patterns left unexplained by simple prey-dependent models, including the apparent stability of food webs and the response of successive trophic levels to ecosystem enrichment (Arditi & Ginzburg, 2012).

Unfortunately the debate remains largely philosophical and based on indirect tests of generic theoretical predictions. Direct empirical evaluations are still limited. These have primarily taken the form of experiments manipulating the abundances of both a predator and a prey (e.g., Fussmann *et al.*, 2005), analyses of predator-prey population dynamics in microcosms (e.g., Jost & Arditi, 2001), and, in rare cases, long-term studies of cooperatively foraging top predator populations (e.g., Vucetich *et al.*, 2002). The majority of these studies have evidenced functional responses that are closer to ratio dependence than to prey dependence (Arditi & Ginzburg, 2012; DeLong & Vasseur, 2011; Skalski & Gilliam, 2001).

Although direct evaluations of predator dependence are increasing in frequency, the logistical and statistical constraints imposed by considering both prey and predator abundances has limited studies to species-poor systems of single predator species interacting with only one primary prey species. Even inherently generalist predators have thereby been reduced to effective specialists, both in manipulative experiments and time-series analyses. Evaluations of the functional forms of interspecific effects between multiple predator species have similarly been inaccessible. Given that most predators in nature are generalists and can alter each others feeding rates in many ways (Kéfi *et al.*, 2012; Peacor & Werner, 2004; Sih *et al.*, 1998) extrapolations regarding the prevalence, strength, and hence importance of predator dependence in species-rich food webs may be premature.

Here we introduce a new approach for characterizing and quantifying the functional responses of generalist predators. By avoiding the logistical constraints imposed by a generalist’s many prey species, the approach may even be used in contexts involving an arbitrary number of interacting predator species. We apply the approach in one set of non-manipulative field surveys and two manipulative field experiments involving two predatory whelks of the Oregon rocky intertidal, *Nucella ostrina* and *N. canaliculata*. Our study of these two predators exposed to their full suite of potential prey provides strong evidence for weak intraspecific predator dependence in *N. ostrina’s* functional response. In the field, over both experimentally-extended and naturally-occurring ranges of predator and prey abundances, this generalist predator is thereby shown to exhibit a functional response that is closer to being prey dependent than ratio dependent. Our study further indicates that *N. ostrina’s* predator dependence is itself prey-specific, with variation in community structure controlling even its qualitative nature. This implies that new functional response models are needed to adequately describe predator-prey interactions in species-rich food webs.

## Methods

We first provide a brief description of the observational approach in order to build intuition for its success. Further details are provided in the *Supplementary Online Materials* (SOM), which also includes descriptions of the functional response models we evaluated in three different contexts. These context (henceforth ‘cases’) were (i) a set of non-manipulative field surveys, (ii) a caging experiment that manipulated predator densities, and (iii) a larger-scale combination of field surveys and predator manipulations, each of which was used to detect or elicit an *in situ* signal of predator dependence.

### The observational approach

Novak & Wootton (2008) introduced a method for inferring the prey-specific per capita attack rates of a generalist predator presumed to exhibit a prey-dependent multispecies type II functional response. Their method is observational in that it uses only data on prey abundances (*N*_*i*_), handling times (*h*_*i*_), and counts of the number of feeding (*n*_*i*_) and non-feeding (*n*_0_) individuals observed during a snapshot survey of a focal predator population. Wolf *et al.* (2015) subsequently showed this method’s analytical estimator for the attack rate on the *i*^*th*^ prey to be equivalent to

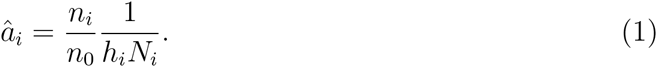

We provide a new and simpler derivation in the SOM.

Intuition for the method’s success may be built by using the attack rate estimator to reformulate the type II functional response model in terms of the fraction of predator individuals that are expected to be observed feeding at any given time. For example, when the predator is a specialist feeding on only one prey species,

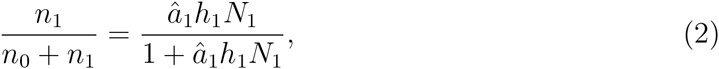

which tends to 1 as *a*_1_, *h*_1_, or *N*_1_ increase. The fraction of individuals observed to be feeding on a particular prey species during a snapshot survey will therefore increase the higher the attack rate, the longer the handling time, or the more abundant the prey species is (Fig. 1A).

**Figure 1:**
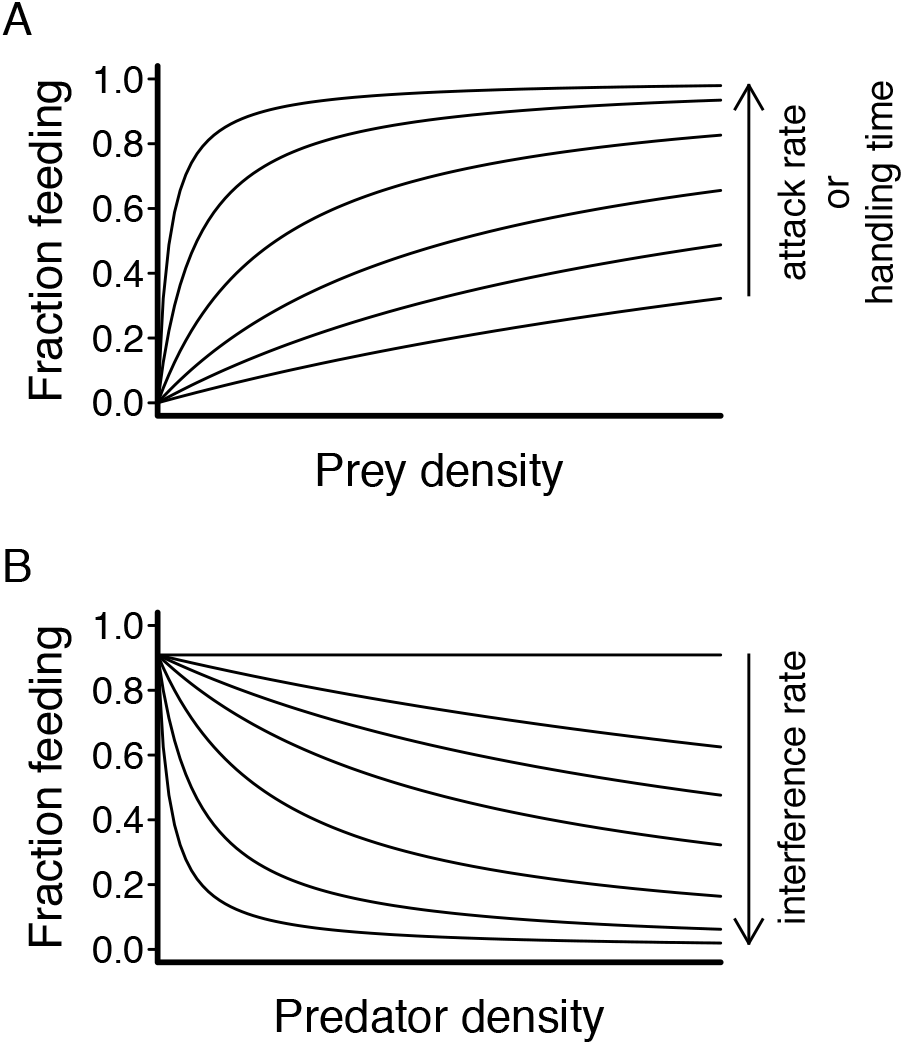
The probability that an individual predator feeding with a type II or Beddington-DeAngelis functional response will be observed in the process of feeding at any point in time (A) increases the higher its attack rate, the longer its handling time, and the more abundant its prey species is (eqn. 2), and (B) decreases with stronger intra- or interspecific interference among predator individuals (eqn. 3). Under the assumption that all individuals are independent and equivalent, this probability corresponds to the fraction of individuals that are expected to be observed feeding in a snapshot survey of the population.

Here we place the Novak & Wootton (2008) method within a general statistical framework, showing eqn. 1 to be the maximum likelihood estimator for the attack rates. This framework enables us to extend the observational approach to situations where ratio-dependent or other, more general, predator-dependent functional response forms are expected, including the Hassell-Varley model (Arditi & Akçakaya, 1990; Hassell & Varley, 1969) and both single- and multi-predator versions of the Beddington-DeAngelis model (Beddington, 1975; DeAngelis *et al.*, 1975). Intuitively, this is possible by considering that the more interference among predators there is, the larger the per capita attack rates must be to maintain the same proportion of feeding individuals. For example, for a specialist predator exhibiting a Beddington-DeAngelis response, the fraction of individuals expected to be feeding at any point in time (Fig. 1B) is described by a binomial likelihood with a probability of ‘success’ equaling

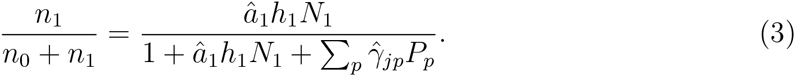

Here *γ*_*jp*_ reflects the per capita strength of the effect of predator species *p* on the focal predator *j*’s feeding rate, and *P*_*p*_ reflects its density. Note that predators can exhibit facilitative effects when *γ* < 0. Correspondingly, the fraction of feeding and non-feeding individuals of a generalist predator population are described by a multinomial likelihood.

Fitting more complex models like the Beddington-DeAngelis model to estimate both the attack rates and mutual predator effects is not possible with only one feeding survey. Rather, doing so requires replicate surveys that vary in predator densities. Specifically, we require at least one more survey than the number of considered predator species. An additional benefit of the statistical framework is that it permits us to evaluate the relative performance of different models in describing empirical data using information theoretics (e.g., AIC). Comparisons can thereby also be made to a simpler (non-functional) density-independent ‘null’ model in which survey-to-survey variation in prey-specific feeding rates is determined not by variation in prey or predator abundances but rather by differences in handling times associated with variation in predator and prey body sizes (see SOM).

### Study system

Our study focused on the species interactions of two intertidal whelks, *Nucella ostrina* and *N. canaliculata*, in midshore ‘mussel-bed patches’. While *N. ostrina* tends to occur higher on the shore than *N. canaliculata*, their tidal range overlaps considerably in the midshore mussel zone where both species often exhibit their highest densities (Connell, 1970; Navarrete, 1996; Spight, 1981). Both species consume the same variety of prey taxa, including mussels, barnacles, limpets, and littorine snails (Palmer, 1984; Spight, 1981). Intertidal whelks like these two species are particularly interesting in the context of functional responses because a key experiment by Katz (1985) involving the Atlantic whelk, *N. lapillus*, has been interpreted by both sides of the debate in support of their arguments (Abrams, 1994; Akçakaya *et al.*, 1995).

Whelk densities are typically highest in patches within the mussel bed where mussels have been removed by wave-induced disturbance (Plate 1; Navarrete, 1996). Patches large enough not to be encroached by the surrounding mussel bed undergo a semi-deterministic trajectory of recovery (Berlow, 1997; Levin & Paine, 1974; Wootton, 2002), being first colonized by diatoms and algae, then acorn barnacles (*Balanus glandula* and some *Chthamalus dalli*), then *Mytilus trossulus* mussels, then *Pollicipes polymerus* gooseneck barnacles, before eventually returning to being dominated by the larger, mussel-bed forming species *Mytilus californianus*. Slow-growing *Semibalanus cariosus* barnacles initiate recruitment in low numbers with the other acorn barnacles but achieve notable densities only at the later stages of succession. At our study site (Yachats, Oregon, 44.3°N, -124.1°W), whelks, limpets (*Lottia asmi, L. digitalis* and *L. pelta*) and littorines (*Littorina sitkana*) are present throughout succession but their abundances vary considerably from patch-to-patch and over time.

### Unmanipulated patches

To quantify attack rates and predator dependence over the natural range of variation in predator and prey densities, we first applied the observational approach to 10 naturally-formed unmanipulated patches. Patches were chosen haphazardly and varied in size (0.8−5.8 *m*^2^, 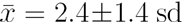) and successional age and thus in species composition, both in terms of absolute and relative species abundances. Species abundances were estimated in each patch using three randomly placed quadrats (25 × 35 *cm*). Low tide feeding surveys were performed in each patch by systematically inspecting and measuring all whelks (±1 *mm*) and noting prey identity and prey size when individuals were feeding (i.e. in the process of drilling, prying or consuming a prey item).

**Plate 1:**
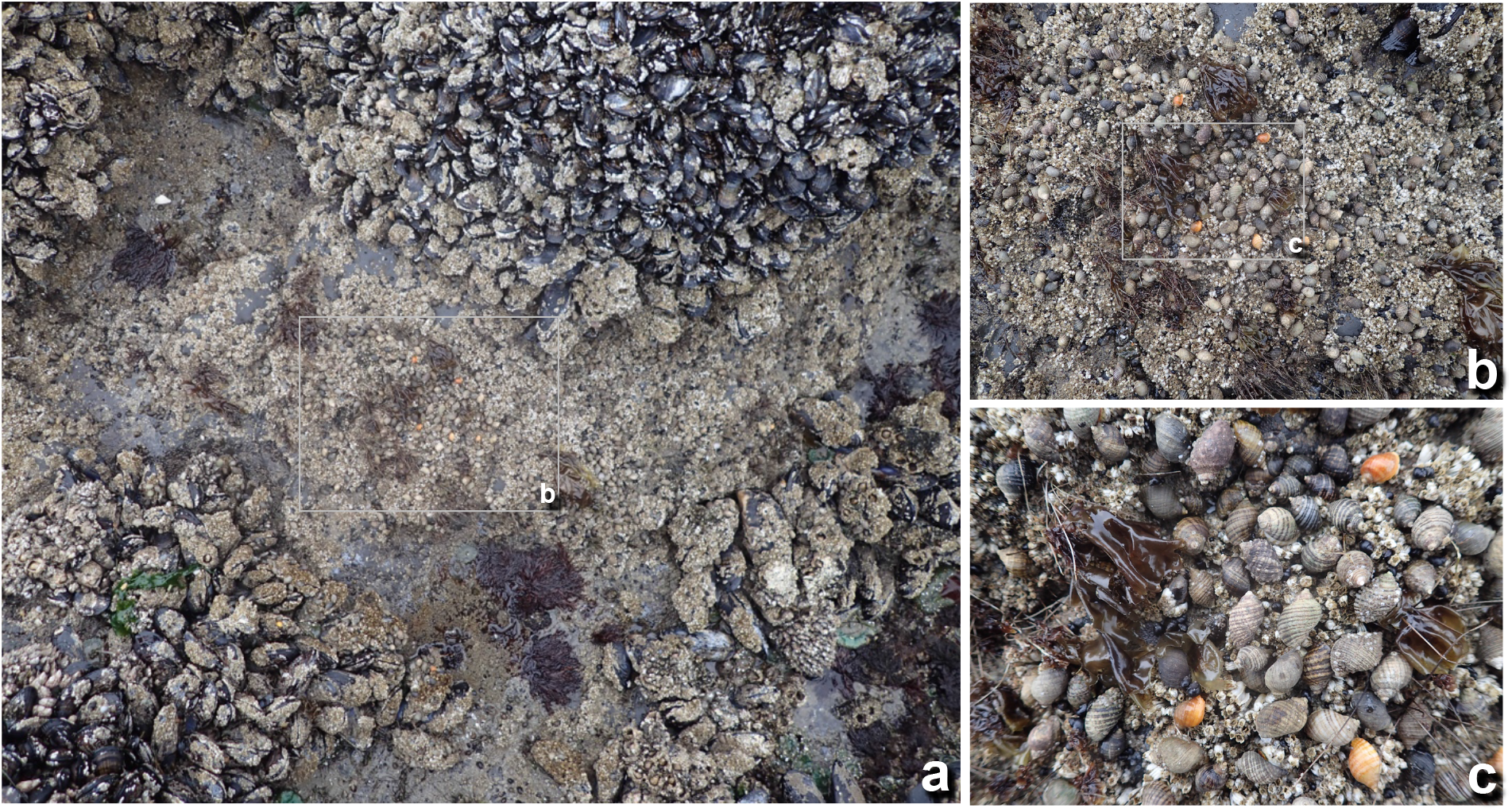
The predatory whelks *Nucella ostrina* and *N. canaliculata* co-occur and can reach extremely high densities in the wave-disturbed patches of a mussel bed.

### Caging experiment

We used a manipulative caging experiment to assess predator dependence over a range of predator densities exceeding that observed in the natural patches, as is typically done in manipulative functional response experiments. Fifteen stainless steel cages (25 × 35 *cm*) were placed in a single large patch of a low-diversity successional age dominated by a homogeneous cover of *Balanus glandula* barnacles. Each cage was photographed to determine prey abundances, then received between 5 and 160 *Nucella ostrina* (11 - 16 *mm* shell length). Feeding surveys of each cage were performed on two subsequent occasions, 2 and 4 weeks later.

### Manipulated patches

Finally, to determine whether predator dependence could be experimentally altered at the patch scale, we combined surveys of naturally-formed patches with a manipulation of their whelk densities. The experiment was performed in 9 haphazardly chosen patches of variable successional age and consisted of five steps: (i) an estimation of species abundances using three haphazardly located quadrats (25 × 35 *cm*); (ii) a first systematic feeding survey of the whelks, (iii) a manipulation of whelk densities, and, after four subsequent high tides, (iv) a re-estimation of whelk densities using three quadrats placed in the same approximate locations as before, and (v) a second systematic feeding survey of the whelks. The manipulation of whelk densities entailed either a decrease or increase (0.07 to 3.3 times their pre-manipulation density), or a control treatment in which all whelks were returned (Fig. S2). Three patches were haphazardly assigned to each treatment. The whelks in all treatments were picked up either during or immediately after the first feeding survey to avoid confounding treatments by the potential effects of whelk handling. Prior observations indicated that a two-day recovery period was ample time for whelks to regain normal activity but insufficient for whelks to have an appreciable effect on prey densities.

### Model-fitting and comparison

Focusing on the feeding observations of *Nucella ostrina,* we estimated the parameters and evaluated the relative performance of five types of multispecies functional response models: type II, ratio-dependent, Beddington-DeAngelis, Hassell-Varley, and the non-functional ‘null’ model. We considered two Beddington-DeAngelis models for the unmanipulated and the manipulated patches in which both *Nucella ostrina* and *N. canaliculata* occurred, one including only an intraspecific predator effect and one including both intra- and interspecific predator effects; only *N. ostrina* was present in the caging experiment.

Our fitting of the models treated the surveys of each case (i.e. the caging experiment, the unmanipulated patches, and the manipulated patches) as independent and identically-distributed, describing feeding rates for each case by one set of attack- and interference rate estimates across all surveys. We also relaxed this assumption for the manipulated patches where two surveys of the same patch had been performed by additionally fitting all models with patch-specific parameters.

Model-fitting involved describing the observed feeding counts by a multinomial distribution whose probabilities were determined by species densities and handling times according to a given functional response model. For the patches, a species’ density was estimated by its mean abundance (*m*^−2^), averaged over replicate quadrats. A species’ handling time was estimated by its mean expected handling time (in hours), averaged over the expected handling times of its feeding observations. The expected handling time of a given feeding observation was estimated from its measurements of whelk- and prey size and ambient temperature (the average of water and air over the month in which surveys were performed) using regression coefficients derived from laboratory experiments manipulating these variables for *Haustrum scobina,* a New Zealand whelk species with ecologically equivalent characteristics and prey (Novak, 2010, 2013). In fitting the models we constrained all attack rates as well as the interference rate parameter of the Hassell-Varley model to be positive. The interference rates of the Beddington-DeAngelis models remained unconstrained. Convergence was reached in all cases by setting the attack rate starting values to reflect the appropriate analytical solutions of the type II or ratio-dependent functional response models (eqns. 1 and S5). Model performance was evaluated by AIC_*c*_ that converges on the AIC goodness-of-fit statistic as sample size increases (Burnham & Anderson, 2004).

## Results

### Variation in diet and species abundances

We observed *Nucella ostrina* feeding on 11 and 10 species, including itself, in the unmanipulated and manipulated patches, respectively. Only 5 of these species were observed being fed upon in the cages, despite the presence of all potential prey and sufficient sampling effort to detect them (Fig. S1). The total number of feeding observations per prey species varied from 2 (*Lotta digitalis*) to 1,089 (*Balanus glandula*), with 14.8% of the 13,131 total examined *N. ostrina* whelks found to be feeding. Six whelks were observed drilling a conspecific individual. *N. ostrina*’s densities ranged between 133 - 1143 *m*^−2^ in the un-manipulated patches, 57 - 1,829 m^−2^ in the cages, and 80 - 1,939 m^−2^ in the manipulated patches prior to manipulation; post-manipulation densities ranged from 80 - 2,518 m^−2^ (Fig. S2). *N. canaliculata*’s densities were consistently and considerably lower (Fig. S2), with only a 128 total feeding observations (14.2% of all examined individuals) being made in the subset of patches in which they were present.

Patches represented early to late successional ages and thus varied considerably in their prey abundances. In particular, the mean densities of *Mytilus trossulus* mussels and *Balanus glandula* barnacles, representing *Nucella ostrina*’s primary prey (both in terms of diet frequency and subsequently estimated feeding rates), respectively varied between 3.8-7,295 m^−2^ and 240 - 114,987 m^−2^. There was no discernible relationship between whelk and prey abundances in the unmanipulated patches (Fig. S3). A positive relationship between *N. ostrina* and *Balanus glandula* densities observed in the manipulated patches prior to manipulation was broken by the manipulation of *N. ostrina* densities (Fig. S3). Patches consequently varied substantially both in the relative ratio of mussels to barnacles and in the relative ratio of whelks to prey (Fig. 2A,C). In contrast, the experimental cages, which were located within a single early successional age barnacle-dominated patch, varied little in their absolute and relative prey abundances (Figs. 2B and S4). The larger-than-natural range of whelk to prey ratios in the cages was therefore due to the manipulation of *N. ostrina* densities. In fitting the alternative functional response models to the data, one prey species, the burrowing mussel *Adula californiensis*, on which two whelks were observed feeding in the unmanipulated patches, was excluded prior to analysis because it was not detected in any abundance survey.

**Figure 2:**
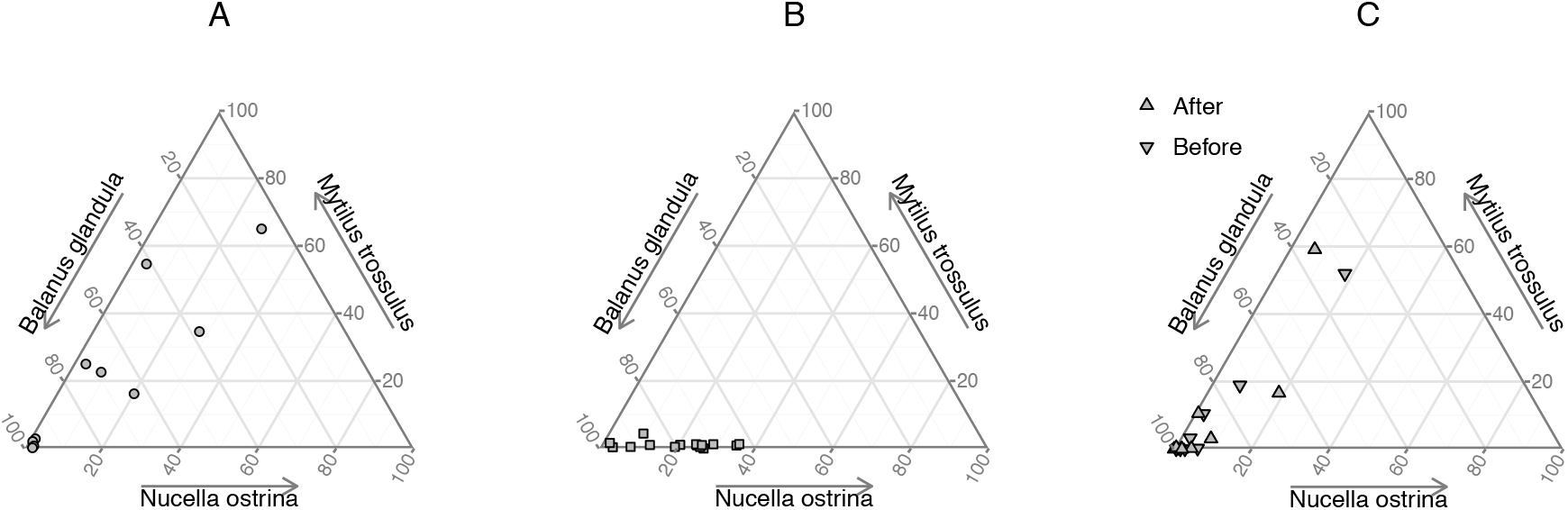
The relative abundance of *Nucella ostrina* and its two primary prey species, *Mytilus trossulus* mussels and *Balanus glandula* acorn barnacles, as illustrated by their proportional densities in the (A) unmanipulated patches, (B) experimental cages, and (C) manipulated patches before and after the manipulation of *N. ostrina*’s densities.

### Model-performance and parameter estimates

The Beddington-DeAngelis functional response entailing only intraspecific predator dependence was unambiguously the best-performing model for the unmanipulated patches; its AIC_*c*_-weight, reflecting the conditional probability of it being the best-performing model, exceeded 0.999 (Table 1A). The patch-specific version of the same model outperformed all others with equally unambiguous evidence for the manipulated patches (Table 1C). Only for the caging experiment did model comparisons fail to provide clear support for a particular model, with the Type II, the Beddington-DeAngelis, and the Hassell-Varley models all exhibiting AIC_*c*_ values within 4 units of each other (Table 1B). Nevertheless, in all three cases the ratio-dependent and density-independent models performed substantially worse than all other models.

**Table 1:**
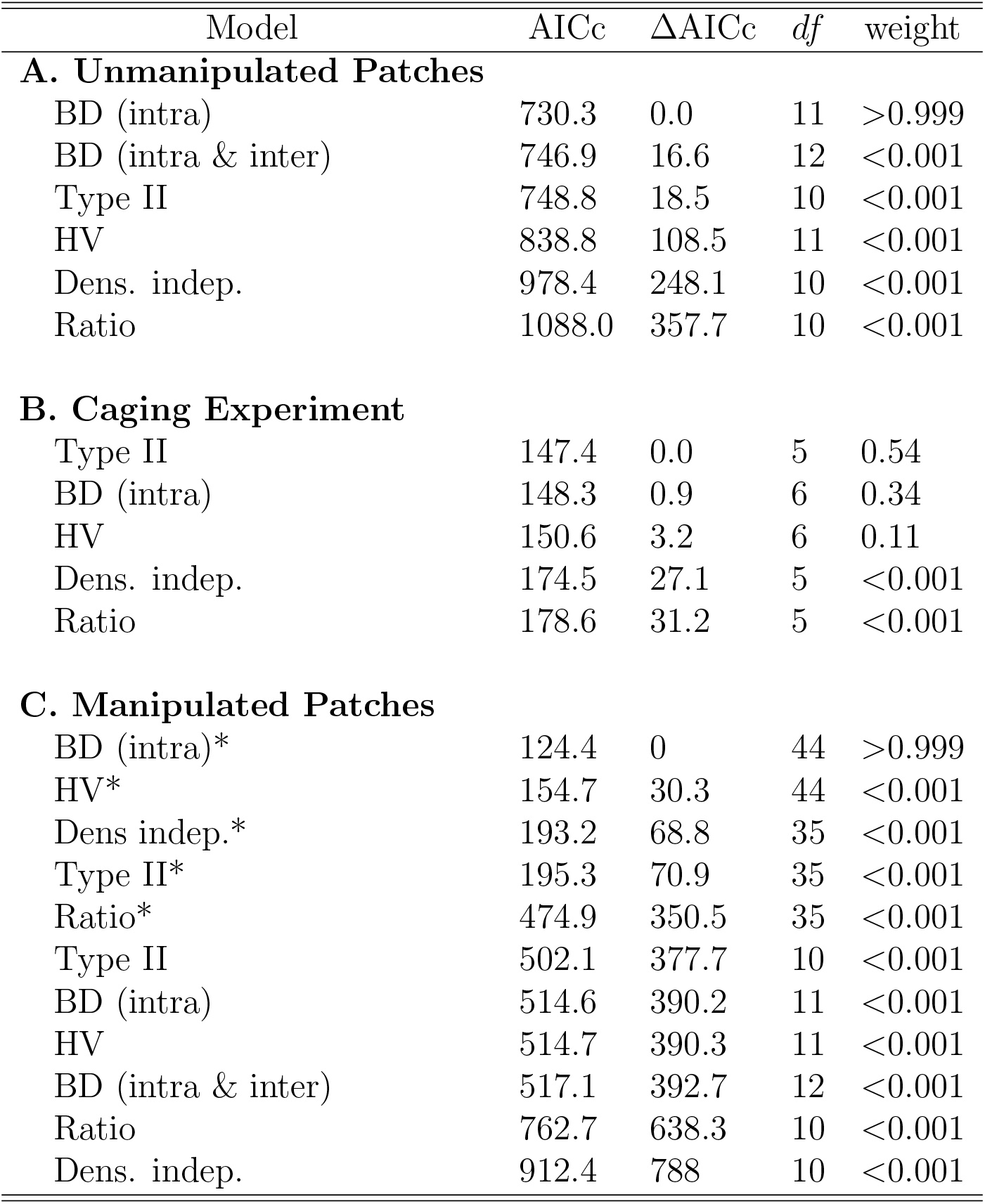
Comparison by AICc of all functional response models applied to (A) the unmanipulated patches, (B) the caging experiment, and (C) the manipulated patches (for which asterisks indicate models with patch-specific parameters). Note that it was not possible to fit the Beddington-DeAngelis model including both intra- and inter-specific effects to the cages or to the manipulated patches on a patch-specific basis.

As estimated assuming the Beddington-DeAngelis model, *Nucella ostrina*’s prey-specific per capita attack rates varied by up to 3 orders-of-magnitude within each of the three cases (Fig. 3A). Attack rates varied over almost 5 orders-of-magnitude across the three cases overall. The range of variation in attack rates was similar in the two sets of patches where *N. ostrina* was observed consuming 10 to 11 species. In the cages, by contrast, the subset of five prey species on which *N. ostrina* was observed feeding evidenced attack rates that were 4 to 1004 times higher than in either set of patches. There was no rank-order correlation between the attack rates of the three cases (Table S10), with a similar number of prey evidencing attack rates that were relatively higher versus lower in one case compared to another. In contrast, although *Nucella ostrina*’s prey-specific feeding rates also varied over 3 orders-of-magnitude, these were of similar magnitude and positively rank-correlated across the three cases (Fig. 3B, Spearman’s *ρ* ≥ 0.7, Table S10).

**Figure 3:**
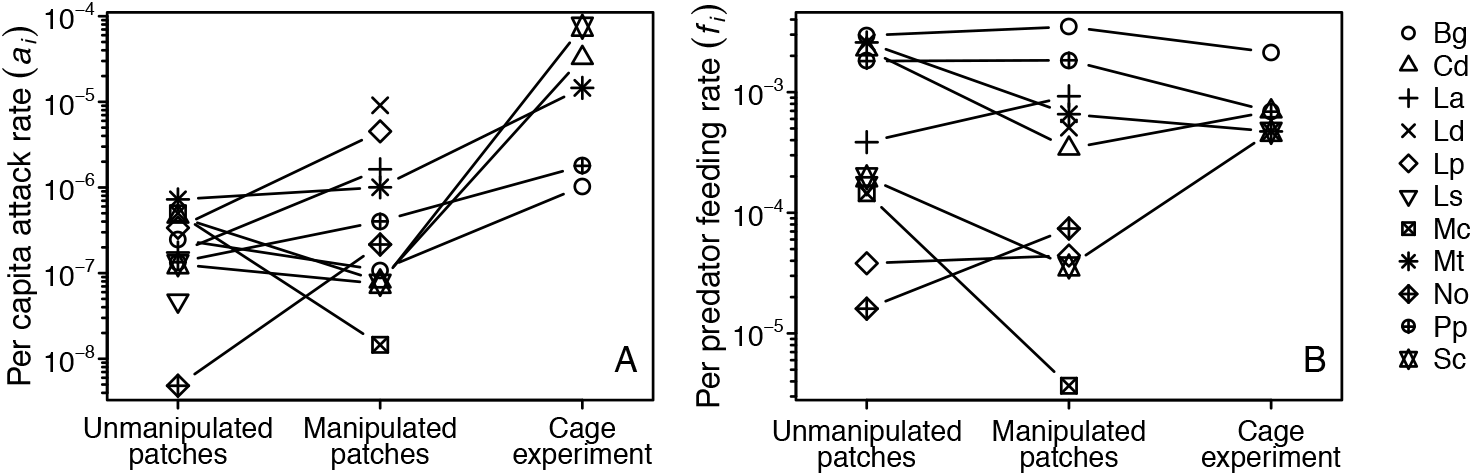
*Nucella ostrina’s* prey-specific per capita attack rates and per predator feeding rates. (A) Per capita attack rate estimates assume a Beddington-DeAngelis functional response with only intraspecific predator effects, and evidence no rank-order correlation between the three cases (Table S10). (B) Feeding rate estimates assume no functional response form and evidence positive rank-order correlations between all pairs of cases (Table S10). Estimates for the manipulated patches are those of the non-patch-specific model. Prey name abbreviations: Bg - *Balanus glandula,* Mt - *Mytilus trossulus;* see Table S1 for others.

Estimates for the per capita magnitude of intraspecific predator dependence in *Nucella ostrina* were larger for the two sets of patches than for the cages (Fig. 4A), consistent with the poorer discrimination among models by AIC_*c*_ for the cages (Table 1). However, while *γ* estimates were positive for both the cages and the manipulated patches (indicating interference effects), the estimate in the unmanipulated patches was negative (indicating a facilitative effect). The patch-specific *γ* estimates for the manipulated patches also exhibited both positive and negative values, with four of the five positive (interference) estimates exhibiting considerably higher magnitudes than the other estimates (Fig. 4B).

**Figure 4:**
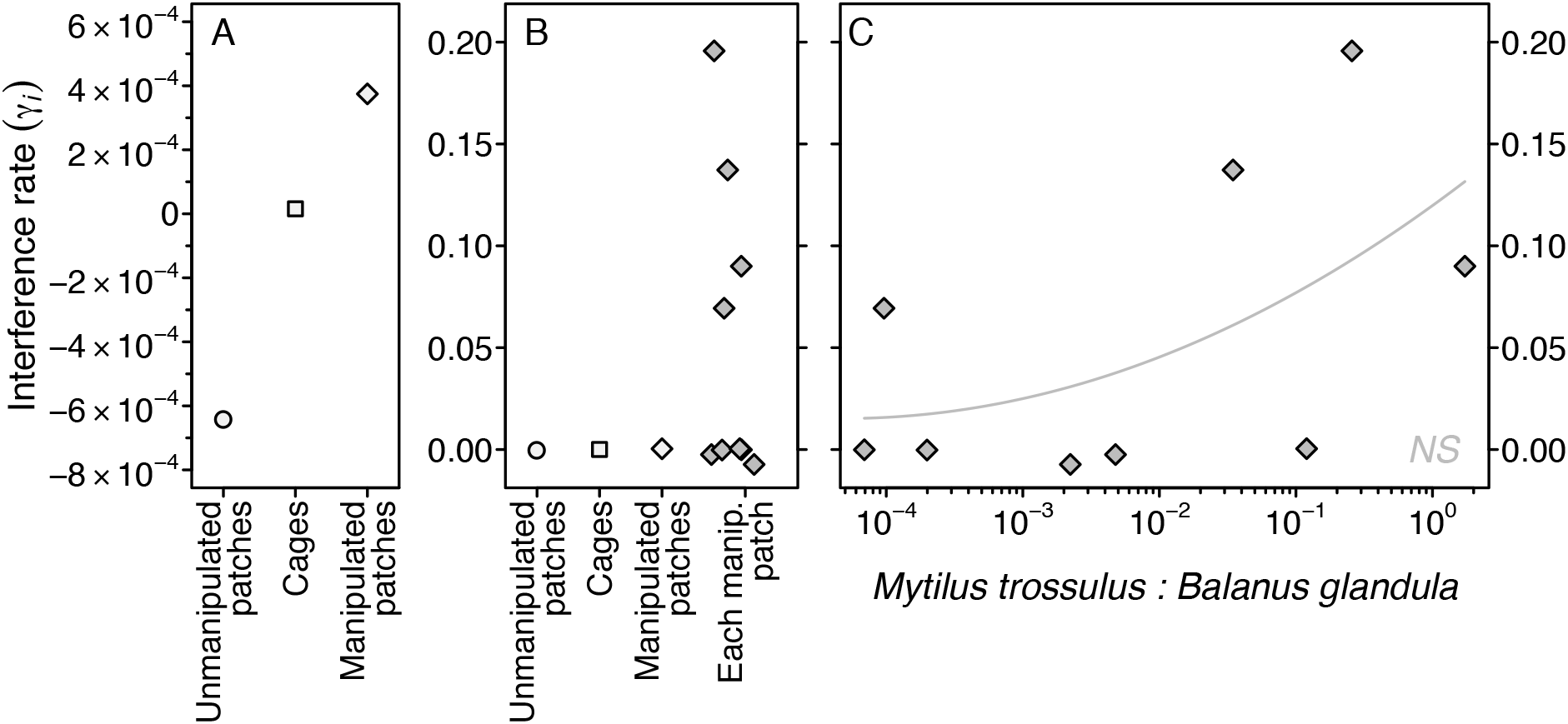
*Nucella ostrina*’s intraspecific predator effects as estimated assuming a Beddington-DeAngelis functional response in (A) each of the three cases (i.e. the unmanip-ulated patches, caging experiment, and manipulated patches) and (B) each case compared to at the patch-specific scale in the manipulated patches. Positive values indicate interference effects while negative values indicate mutualistic effects. (C) Patch-specific predator effect estimates versus the relative density of *N. ostrina*’s two primary prey, *Mytilus trossulus* mussels and *Balanus glandula* barnacles, in the manipulated patches suggests that the per capita strength of predator interference may depend on prey abundances. The fitted second-order polynomial trendline is *not* significant (*R*^2^ = 0.3, *p* = 0.34). Note different y-axis scales in A versus B and C.

## Discussion

Two fundamental yet often conflated questions have contributed to sustaining the debate over predator functional responses: How to best represent predator-prey interactions in models of population dynamics? And, what are the mechanistic relationships between predator feeding rates and species abundances? The importance of these questions transcends predator-prey interactions (Abrams, 2015; Perretti *et al.*, 2013). Indeed, all methods for quantifying the strengths and hence importance of species interactions make assumptions regarding their functional form (Novak *et al.*, 2016; Vázquez *et al.*, 2015; Wootton & Emmerson, 2005). Recognizing that the answers to these two questions may not be the same will be key to future progress. For example, predator dependence may be sufficiently weak that it has no appreciable effect over the range of species abundances that actually occur in nature’s species-rich communities, despite being discernible in manipulative experiments (Fussmann *et al.*, 2005).

### Strong evidence for weak predator dependence

Our study indicates that predator dependence as encapsulated by Beddington-DeAngelis model characterizes *Nucella ostrina*’s functional response the best, and that its effects are discernible over the species abundances and diversity of prey that this generalist predator experiences in the field. The per capita strength of predator dependence was nonetheless weak. This was most clearly evidenced by the relative performance of the next-best prey-dependent type II model (Table 1), and by the point estimates for the interference-strength parameter of the Hassell-Varley model (*m* ≤ 3.28 · 10^−5^ in all three cases, where *m* = 0 reflects complete prey dependence, Tables S2 - S4). In contrast, the ratio-dependent model, which has seen more theoretical treatment than any other predator-dependent model (Arditi & Ginzburg, 2012), was consistently among the worst-performing models. In two of three cases it performed even more poorly than the model which assumed feeding rates to be independent of species abundances altogether (Table 1). Our study thereby distinguishes itself in that most previous studies have dichotomously concluded functional responses to be best described by either a prey-dependent or the ratio-dependent model, despite general recognition that the truth is somewhere in between (Abrams, 2015; Abrams & Ginzburg, 2000; Arditi & Ginzburg, 2012).

Surprisingly, our analysis inferred no effect of *Nucella canaliculata* on *N. ostrina*’s feeding rates, despite their seeming ecological similarities. This may have been due to insufficient statistical power associated with low replication (*n* ≤ 10 patches) and the relatively low variation seen in *N. canaliculata’s* abundances (Fig. S2); the Beddington-DeAngelis model including both inter- and intraspecific predator effects did perform best in the two sets of patches when model performance was evaluated by AIC rather than AIC_*c*_ (Table S11). However, an implicit benefit of the observational framework is that its focus on the fraction of feeding individuals makes it most sensitive to the effects of predator dependence at low predator densities, where a doubling of predator numbers has a larger effect on per individual feeding rates than it does at high predator densities (Fig. 1B). This differs from the approach of traditional functional response experiments where the largest and thus most easily estimated rates of overall prey removal occur at high predator densities. Therefore, *N. canaliculata’s* low densities should not have been an issue. Our results therefore suggest that the interaction between the two whelk species is primarily one of indirect effects mediated by prey exploitation, rather than representing a significant interaction modification of feeding rates (Kéfi *et al.*, 2012; Spight, 1981).

Similarly unexpected was that the weakest support for predator dependence was seen in the caging experiment where its effects were most expected (Table 1); the experiment manipulated *N. ostrina’s* densities beyond their typical range and affected predator-prey ratios exceeding their natural variation (Figs. 2 and S2). Furthermore, the prey depletion that likely occurred between the initiation of the experiment and when the feeding surveys were conducted should have favored predator-dependent models by reducing feeding rates most in the high density cages.

One explanation for the experiment’s inability to discriminate among models more clearly was that the fraction of feeding individuals will not have been estimated as reliably in the low predator density cages. Given the dimensions of a cage, the number of whelks in the lowest density cage was only five, for example. Thus the probability of observing all or none of the individuals feeding at any given time was high regardless of their true mean feeding rate. This issue will have been alleviated by our use of repeated cage surveys (*n* = 30 surveys), and was altogether avoided for the much larger natural patches that each contained many more whelks in total.

A more likely explanation for the weak expression of predator dependence in the caging experiment is that the cages, or their placement within an early successional age patch that was dominated by a single barnacle species, altered whelk foraging behavior from that exhibited across the sets of surveyed patches more generally. This interpretation challenges the concern that such more traditional functional response experiments involving isolated predator-prey pairs could be favoring the detection of predator dependence by selecting for and magnifying the strength of strong predator-prey interactions (Abrams, 2015). However, the results of our analysis are also consistent with this concern in that *Nucella ostrina*’s per capita attack rates were substantially higher in the cages (Fig. 3A). Indeed, the observation that the highest prey-specific feeding rates decreased while the lowest prey-specific feeding rates increased in the cages relative to the patches, even as their overall rank-order remained relatively consistent across the three cases (Fig. 3B), suggests that the caged whelks altered their foraging strategy to compensate for the reduced breadth of their diet.

### Prey-specific predator dependence

While further experiments involving generalist predators will be needed to determine how diet breadth itself can affect the strength of predator dependence, a likely feature distinguishing the functional responses of generalist and specialist predators is the variable propensity of a generalist’s different prey species to elicit predator dependence. For whelks in particular, predator dependence will have been driven by a number of mechanisms that vary by prey identity and differ in their qualitative nature.

For example, two mechanisms of interference that we observed directly were the drilling of conspecific individuals and the simultaneous feeding on the same prey item by two individuals. Similar mechanisms of negative predator dependence are commonly invoked in the literature (Arditi & Ginzburg, 2012). Conspecific drilling represents time wasted in regards to further foraging opportunities, even when consumption itself does not occur. Its frequency would typically be expected to increase with predator density irrespective of prey identity, but this was not observed in our study (Fig. S5). Extensive surveys at a nearby study site nonetheless show that the shells of least 0.1% of the *N. ostrina* population bear the mark of drilling events (MN, *unpubl. data*). In turn, the simultaneous feeding by two individuals on the same prey item represents reduced energetic payoff, which may also be substantial for whelks given their long handling times. In contrast to conspecific drilling, we observed simultaneous feeding almost exclusively when whelks fed on *Mytilus trossulus* mussels, a likely consequence of the large surface area for drilling that a mussel shell represents, the longer handling time of the average mussel relative to other species, and the tendency of mussels to form clusters around whose accessible perimeters whelk densities are often locally increased (see also Hossie & Murray, 2016).

Much less considered in the debate over functional responses is that predator density can also have positive facilitative effects on feeding rates, even in the absence of cooperative group hunting. This omission persists despite the longstanding awareness of the synergistic effects between predator species (Sih *et al.*, 1998). For whelks, a potentially common mechanism for facilitative effects is the feeding-induced release of prey chemical cues. That this mechanism can be prey-specific has recently been demonstrated by the characterization of a cuticular glycoprotein in *Balanus glandula* that acts as a potent stimulant for whelk feeding, the nature of which is specific to acorn barnacles (Zimmer *et al.*, 2016).

If both facilitative and interference-based mechanisms of predator dependence exist and are dependent upon prey identity, then, for generalists, both the strength and net qualitative nature of predator dependence should depend on community structure. This appears to have been the case in our study, with *γ* estimates for the Beddington-DeAngelis model indicating (1) net interference in the manipulated patches where *Mytilus trossulus* mussels tended to be more common, (2) weaker interference in the cages where *Balanus glandula* barnacles were dominant, and (3) net facilitation in the unmanipulated patches where a second barnacle species tended to be more common (Table 1, Figs. 2 and S4). Further support is suggested by our patch-specific analysis of the manipulated patches, with *γ* estimates tending to increase with the ratio of available mussels and barnacles (Fig. 4C). Future experiments manipulating community structure directly will be needed to determine whether such prey-specific influences of community structure tend to be idiosyncratic or conform to useful categorizations.

## Conclusions

That many prey-specific mechanisms of predator dependence are likely to occur in the functional responses of generalist predators indicates that additional, more complex models will be useful in characterizing the species interactions of nature’s species-rich food webs. Many more such models, including those that relax the assumptions of predator homogeneity and the constancy of per capita rates (e.g., Baudrot *et al.*, 2016; Chesson, 1984; Murdoch & Oaten, 1975), should become empirically accessible with the observational framework, particularly when applied in combination with experimental manipulations. Additional matters of ‘instantism’ in the parameterization of dynamical population models that have plagued the interpretation of traditional functional response and interaction strength experiments (see Fussmann *et al.*, 2007; Jensen *et al.*, 2007; Novak & Wootton, 2010) should also be assuaged by the now logistically feasible repeated application of the observational approach over the biologically appropriate time-scales of a focal predator’s numerical response.

## Acknowledgments

We are grateful to Shannon Hennessey, Stephanie Merhoff and Beatriz Werber for their assistance performing the manipulated patch experiment, and to Ben Dalziel and Dan Preston for comments on the manuscript. Method development was supported in part by NSF DEB-1353827. Data and example code are available on the Dryad Digital Repository (doi: *post-acceptance*) and at https://github.com/marknovak/PredDep.

